# Direct RNA Sequencing of the Complete Influenza A Virus Genome

**DOI:** 10.1101/300384

**Authors:** Matthew W. Keller, Benjamin L. Rambo-Martin, Malania M. Wilson, Callie A. Ridenour, Samuel S. Shepard, Thomas J. Stark, Elizabeth B. Neuhaus, Vivien G. Dugan, David E. Wentworth, John R. Barnes

## Abstract

For the first time, a complete genome of an RNA virus has been sequenced in its original form. Previously, RNA was sequenced by the chemical degradation of radiolabelled RNA, a difficult method that produced only short sequences. Instead, RNA has usually been sequenced indirectly by copying it into cDNA, which is often amplified to dsDNA by PCR and subsequently analyzed using a variety of DNA sequencing methods. We designed an adapter to short highly conserved termini of the influenza virus genome to target the (-) sense RNA into a protein nanopore on the Oxford Nanopore MinION sequencing platform. Utilizing this method and total RNA extracted from the allantoic fluid of infected chicken eggs, we demonstrate successful sequencing of the complete influenza virus genome with 100% nucleotide coverage, 99% consensus identity, and 99% of reads mapped to influenza. By utilizing the same methodology we can redesign the adapter in order to expand the targets to include viral mRNA and (+) sense cRNA, which are essential to the viral life cycle. This has the potential to identify and quantify splice variants and base modifications, which are not practically measurable with current methods.

## Introduction

Decades ago, a method was published describing the use of base-specific chemical degradation with chromatographic and autoradiographic resolution as a way of directly sequencing short stretches of RNA^1^. Since then, little progress has been made on directly sequencing RNA. Instead, the elucidation of RNA sequences is typically indirect and primarily requires methods that synthesize cDNA from RNA templates. While these methods are powerful^2^, they suffer from limitations inherent to cDNA synthesis and amplification such as template switching^3^, artifactual splicing^4^, loss of strandedness information^5^, obscuring of base modifications^6^, and propagation of error^7^. In 2009, a method for RNA sequencing was developed on the Helicos Genetic Analysis System where poly(A) mRNA is sequenced by the step-wise synthesis and imaging of nucleotides labeled with an interfering but cleavable fluorescent dye^8^. While the input material requirements for this method are extremely low, the long workflow and short reads are limiting. Nevertheless, these approaches expose two major limitations of RNA sequencing: sequencing by synthesis and short read length. Overall, current technologies for sequencing RNA templates present difficulties in the assessment of base modifications, splice variants, and analysis of single RNA molecules.

Influenza viruses are negative-sense segmented RNA viruses^9–11^, and sequencing these viruses has played an important role in their understanding for 40 years^12,13^ including the discovery of highly conserved viral RNA termini^14^ (**Figure 1A**). These 3’ and 5’ termini are 12 and 13 nucleotides in length, respectively, and they are highly conserved across the PB2, PB1, PA, HA, NP, NA, M, and NS genome segments of influenza A viruses, which enabled the development of a universal primer set for influenza A virus genome amplification^15,16^. Even though these conserved vRNA termini have been readily exploited for efficient next generation sequencing (NGS) of influenza virus segments^16–18^, current methods retain some of the limitations inherent to cDNA-based techniques^3–7^. A new tool for long read direct RNA sequencing could reduce these biases and greatly aid efforts to directly sequence influenza virus and other RNA viruses.

Oxford Nanopore Technologies (ONT) recently released their direct RNA sequencing protocol. This method involves the sequential ligations of a reverse transcriptase adapter (RTA) and a sequencing adapter^19^. The RTA is a small dsDNA molecule (**Figure 1B**) that contains a T10 overhang designed to hybridize with poly(A) mRNA and a 5’ phosphate (Pi) that ligates to the RNA creating a DNA-RNA hybrid. The RTA also serves as a priming location for reverse transcription of the entire length of the RNA molecule, though the cDNA generated is not sequenced. The DNA-RNA hybrid is then ligated to the sequencing adapter which directs the RNA strand of the assembled library into the nanopore for sequencing^19^.

**Figure 1.**
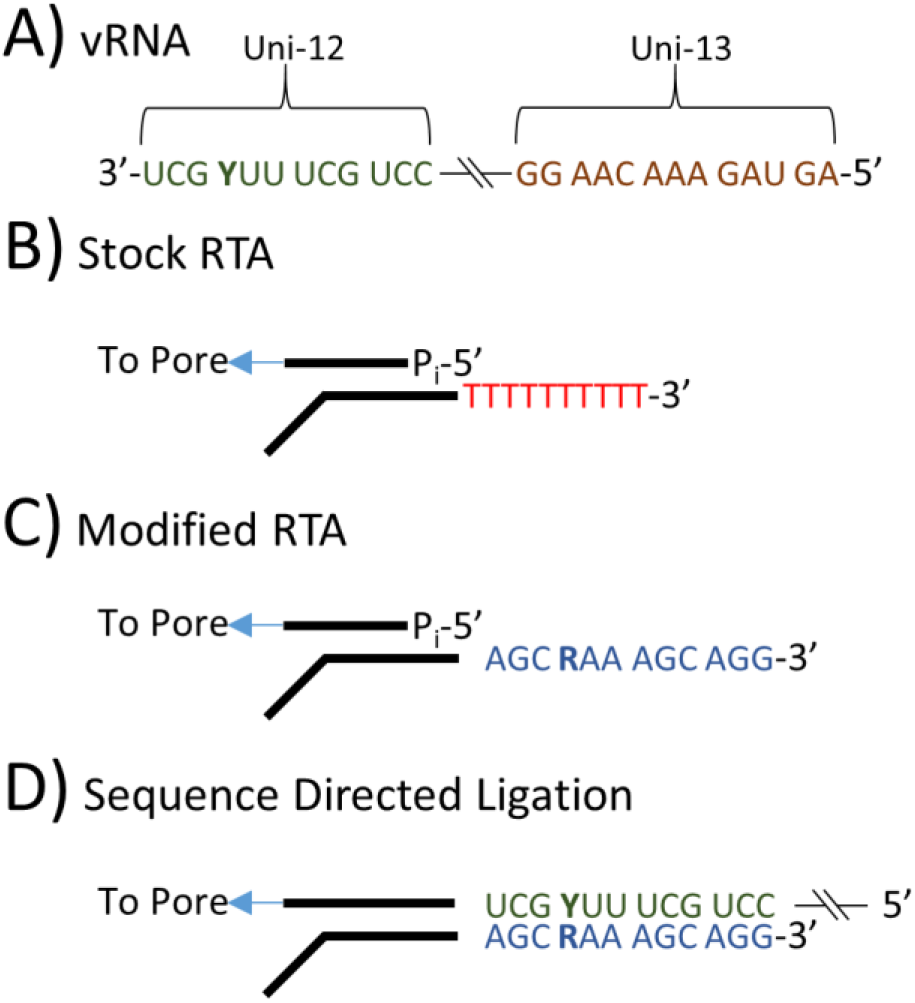
A) Influenza A viruses contain highly conserved 12 and 13 nt sequences at the 3’ and 5’ termini. B) The key component of Oxford Nanopore direct RNA sequencing is a Reverse Transcriptase Adapter (RTA) which targets poly(A) mRNA and is ligated to the 3’ end of the mRNA. A sequencing adapter is then ligated to the RTA which directs the RNA strand into the pore for sequencing. C) The RTA was modified to target the 3’ conserved 12 nt of the influenza A virus genome. D) The modified RTA hybridizes and is ligated to vRNA in the first step of direct RNA sequencing.

We describe direct RNA sequencing of an influenza A virus genome through modification of recently released RNA methods from Oxford Nanopore Technologies^19^ (**Figure 1C**) by targeting the conserved 3’ end of the genome with an adapter to capture it (**Figure 1D**), rather than a primer to amplify it. The efficacy of the adapter is tested by sequencing the RNA genome of an influenza virus generated by reverse genetics A/Puerto Rico/8/1934 (PRJNA449380). The RNA was isolated from virus containing allantoic fluid (crude) harvested from infected embryonated chicken eggs. The results from the nanopore sequencing are compared to the current Illumina-based pipeline utilized by the Influenza Genomics Team at the Centers for Disease Control and Prevention.

## Results

### Nanopore sequencing

First, the RNA calibration strand enolase was directly sequenced on the MinION platform. Three sequencing experiments covered 100% of the 1,314 nucleotide long RNA molecule to an average depth of 122,207 ± 8,126 (sd). Of the 169,041 ± 28,741 reads, 98.6 ± 1.7% mapped to the reference sequence (**Table 1**), with 100% of the mapped reads in the sense orientation. The direction of the reads and the positive slope of the coverage diagram (**Figure S1**) are indicative of directional sequencing of mRNA from the 3’ end. The distribution of read lengths (**Figure S2 and Table S1**) accurately corresponds to the expected length of 1,314 nucleotides. The read level accuracy was 90.4 ± 0.8%, and the consensus sequence was 99.7% in concordance with the known reference.

**Table 1.** Individual MiSeq experiment is compared to MinION experiments of enolase mRNA (technical triplicate) and influenza vRNA from crude virus (triplicate). Values from triplicate experiments are presented as averages ± standard deviation.

|  | MiSeq<br>Crude | MinION |  |
| --- | --- | --- | --- |
|  |  | Enolase | Crude |
| Reads | 143,572 | 171,135 ± 26,929 | 54,353 ± 15,314 |
| Mapped Reads | 143,378 | 169,041 ± 28,741 | 53,721 ± 15,145 |
| % Mapped | 99.9% | 98.6 ± 1.7% | 98.8 ± 0.1% |
| Accuracy | 99.6% | 90.4 ± 0.8% | 86.2 ± 0.31% |
| Insertion | 0.30% | 1.49 ± 0.02% | 1.66 ± 0.01% |
| Deletion | 0.06% | 5.4 ± 0.5% | 8.2 ± 0.2% |
| Substitution | 0.36% | 4.7 ± 0.4% | 6.4 ± 0.1% |
| Consensus | ≡ 100% | 99.7% ± 0% | 98.97 ± 0.01% |

Based on available details on the RTA system, it was possible to make further modification to target other RNA species (**Figure 1**). To adapt this technique for the influenza virus genome, the target sequence of the RTA was changed from an oligo-dT to a sequence complementary to the 12 nucleotides that are conserved at the 3’ end of the RNA segments of influenza A viruses (**Table S2**).

To test the effectiveness of the modified adapter, total RNA from allantoic fluid (crude) harvested from infected chicken eggs was sequenced via MinION. Three sequencing experiments covered 100% of the PB2, PB1, PA, HA, NP, NA, M, and NS gene segments to an average depth of 3,269 ± 1,892 (**Figure 2**). Although, there is reduced coverage at the extreme termini (**Figure 3**) and a heavy coverage bias towards the 3’ terminus of the negative sense RNA, since this approach reads from the 3’ to 5’ end of the molecule. Of the 54,353 ± 15,314 reads, 98.8 ± 0.1% mapped to influenza (**Table 1**) in a roughly even distribution among the 8 segments (**Figure S3**), with 100% of the mapped reads in the negative-sense orientation. The distribution of read lengths (**Figure 4 and Table S1**) corresponds well to the expected length of the respective segment. The read level accuracy was 86.3 ± 0.3%, and the consensus sequence was 98.97 ± 0.01% in concordance with consensus sequence generated using our modified version of the multi-segment reverse transcriptase polymerase chain reaction (M-RTPCR)^15,16^, Nextera, and MiSeq approach.

**Figure 2.**
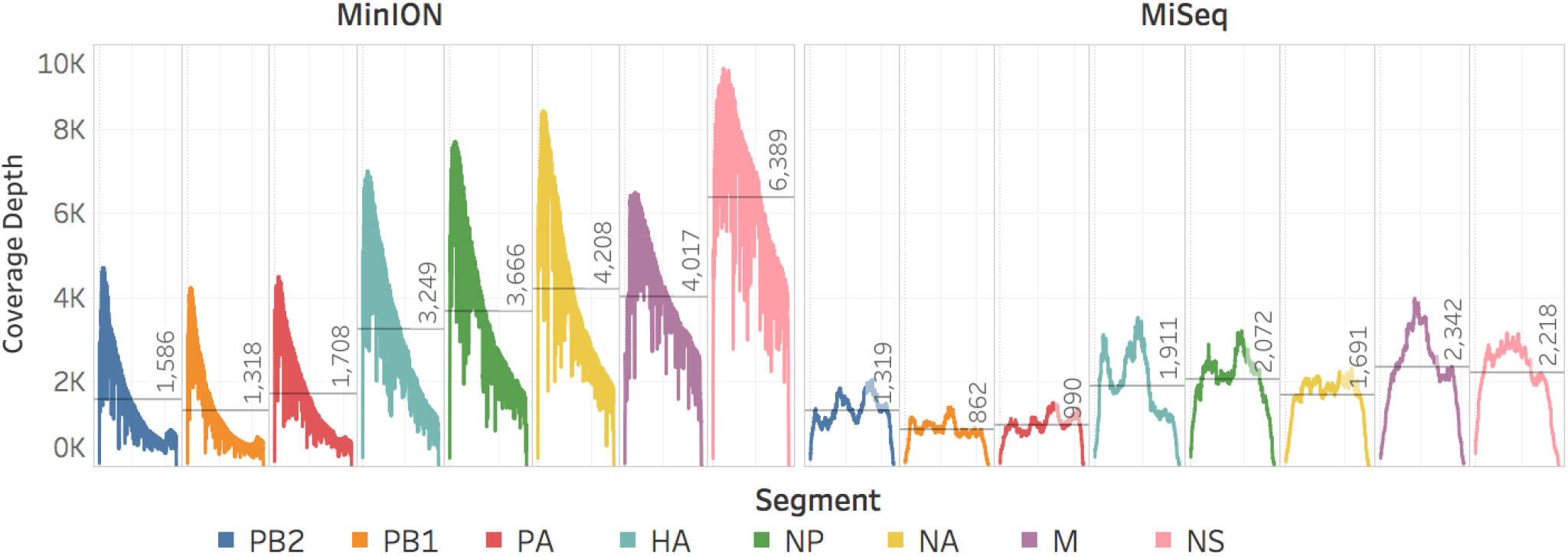
MinION and MiSeq sequencing coverage of the PB2, PB1, PA, HA, NP, NA, M, and NS genome segments of the influenza virus genome from the crude viral samples. Negative-sense slope coverages in the MinION results confirm the directionality of the sequencing and capture method.

**Figure 3.**
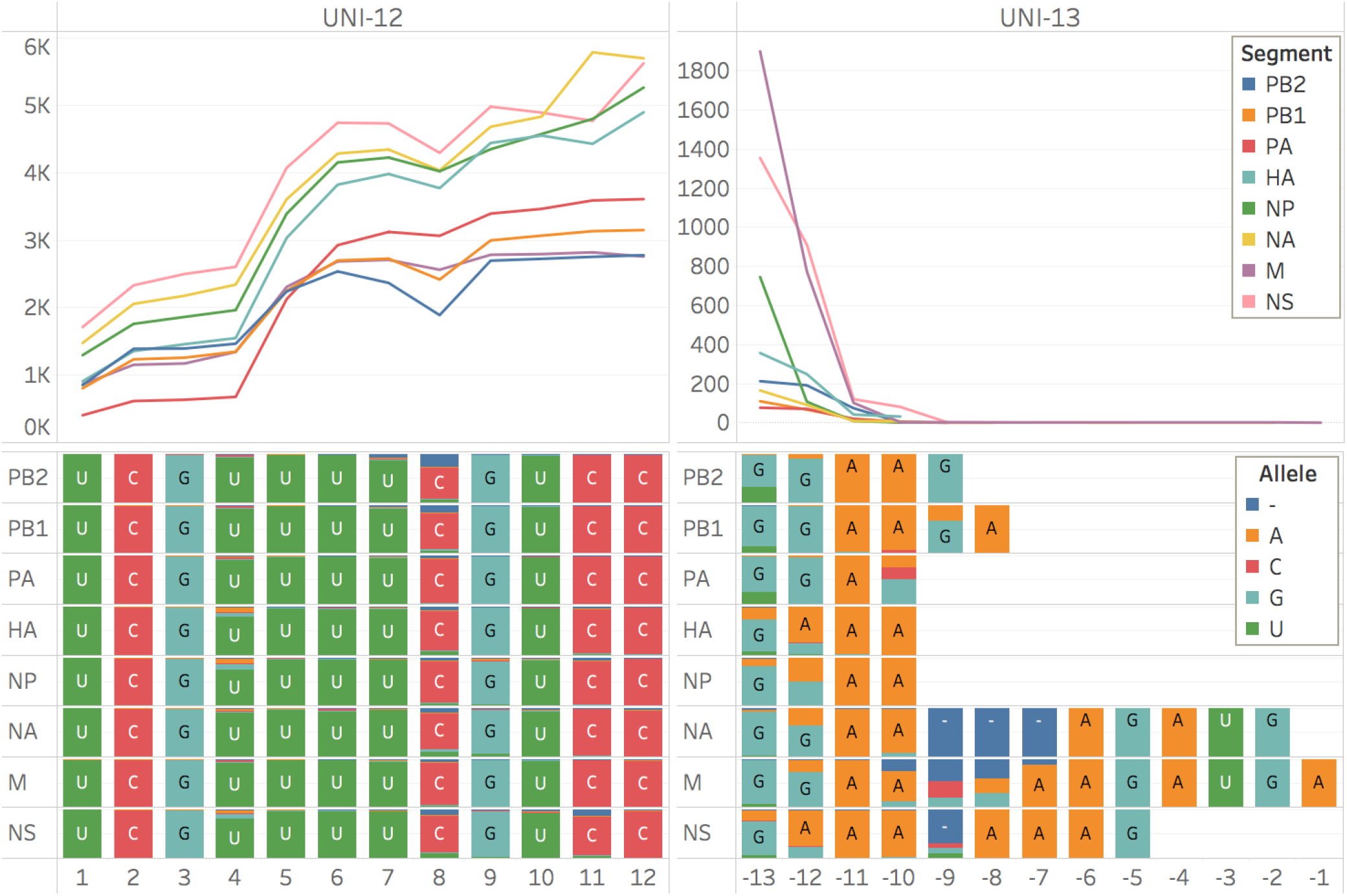
The aligned read length distributions correspond to the expected lengths (dashed lines) of the respective segments (NS 890 nt; M 1,027 nt; NA 1,413 nt; NP 1,565 nt; HA 1,778 nt; PA 2,233 nt; PB1 and PB2 2,341 nt) from the crude viral samples. As the segment length increases, the read length distribution falls further short of the expected length, presumably due to RNA degradation. Aligned read lengths include insertion errors, accounting for the presence of reads larger than the expected value. Due to cases of large insertion errors, 14 total reads longer than 2,500 nucleotides were observed.

**Figure 4.**
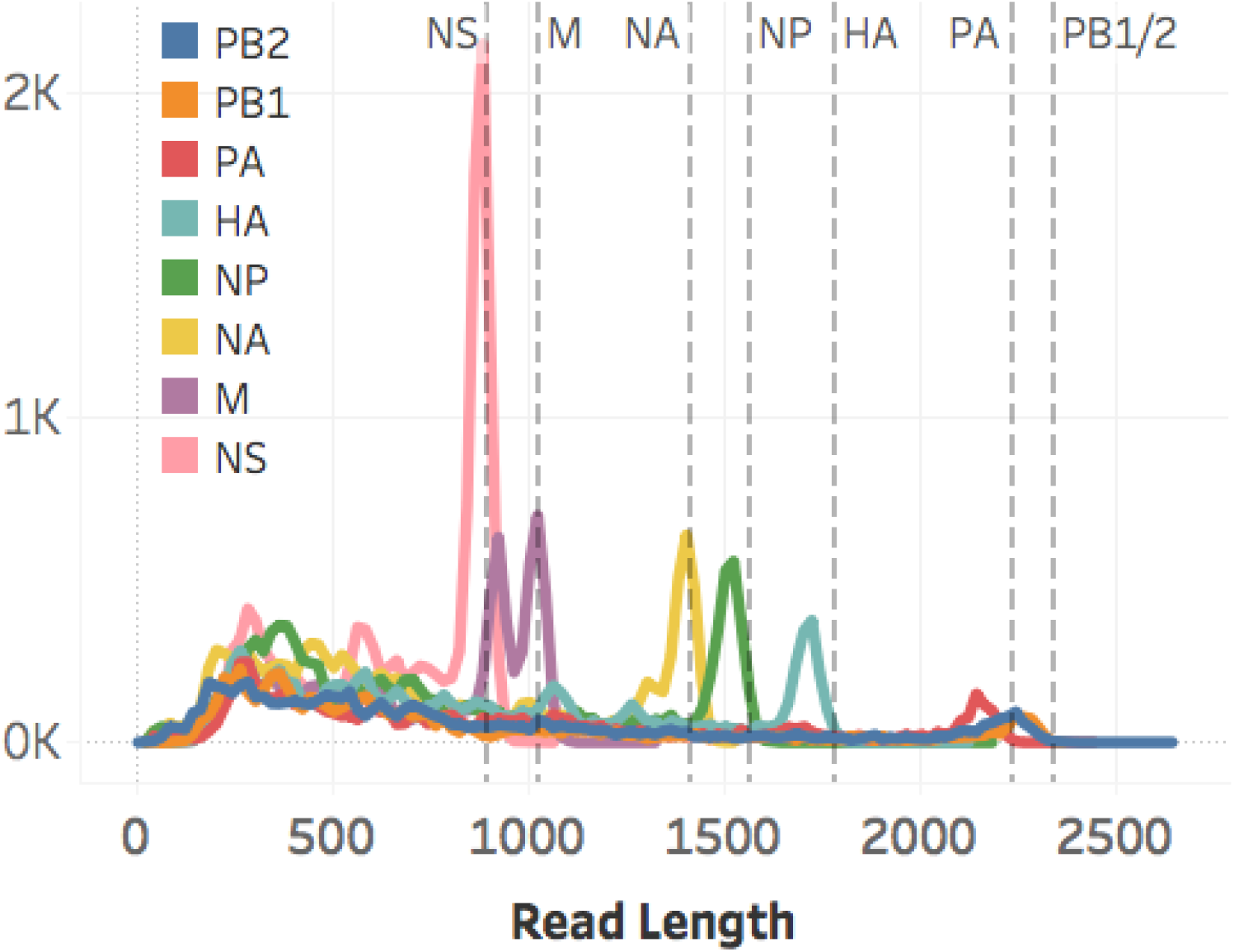
MinION coverage and consensus of the conserved 3’ (Uni-12) and 5’ (Uni-13) termini of the influenza virus RNA segments from the crude viral samples.

To provide a favorable substrate for the modified adapter and a positive control for future experiments, RNA from two sucrose purified virus preparations (pure) was sequenced via MinION. Two sequencing experiments covered 100% of the PB2, PB1, PA, HA, NP, NA, M, and NS viral RNA segments to an average depth 9,312 and 1,068 respectively (**Figure S4**). Of the 119,860 and 13,848 reads acquired in each run, 99.6 and 99.1% mapped to influenza virus (recombinant A/Puerto Rico/8/1934), respectively (**Table S3**), in a roughly even distribution among the eight vRNA segments (**Figure S3**) with 100% of the mapped reads in the negative-sense orientation. The distribution of read lengths (**Figure S5 and Table S1**) corresponds to expected lengths of each respective segment. The read level accuracies for the two runs were 85.2 and 83.8%, and the consensus sequences were 98.7 and 98.5% in concordance with consensus sequence generated using our standardized M-RTPCR amplified genome and MiSeq approach.

### Illumina MiSeq sequencing

The viral RNA segments from the pure and crude preparation were amplified by M-RTPCR^15,16^, and size fractionation of those amplicons showed the characteristic banding pattern of the amplified influenza virus genome (**Figure S6**). Sequencing of the RNA from purified virus or crude virus produced 163,264 and 143,572 reads, respectively, of which 99.9% mapped to influenza A virus (**Table 1**). The reads were roughly evenly distributed among the 8 segments (**Figures S3**). The mapped reads covered 100% of all 8 genome segments (**Figures 2 and S4**) with reduced coverage at the extreme termini (**Figure S7**). The read level accuracy was 99.6% and the consensus sequences, which were used as the reference genome for the nanopore assemblies, were defined as 100% accurate and were 100% identical to each other.

## Discussion

We have demonstrated, for the first time, complete^20^ sequencing of an RNA virus genome by direct RNA sequencing. Using a method originally designed to sequence mRNA, we adapted the target sequence to bind the 3’ sequence conserved among influenza A viruses. The specificity of this adapter allowed efficient sequencing of influenza virus RNA genomic segments from RNA isolated from purified virus particles (control) or from RNA isolated from a crude extract that contains a myriad of viral and host (chicken) RNAs. Using this adapter, 98.8% of reads from the crude RNA preparation mapped to the influenza virus, which is practically as efficient as with purified virus RNA sample (99.3%). This performance on crude virus stocks demonstrates that the sequence directed library preparation is a very effective method to select specific target RNA species among a population of RNAs, as the vast majority of reads were to A/Puerto Rico/8/1934 using 12 ribonucleotides as the target sequence.

The data shows that other modifications to the adapter could target other RNA species such as RNAs from specific pathogens and different RNA species within a particular pathogen. For example, one could compare (+) sense cRNA [replication intermediate of (-) sense vRNAs], (+) sense mRNAs, or (-) sense RNAs present during RNA virus infections (such as for influenza viruses). The data illustrates that the adapter sequence could be modified to target specific viral families, genus, or species by extending the target sequence and or by adding degeneracies. This is an advantage over poly(A) methods that have a reduced signal-to-noise ratio due to host mRNA. Targeting influenza A virus vRNA and cRNA independently may prove difficult as there is complementarity between the two conserved termini of the vRNA segments, and therefore high sequence identity between the 3’ termini of the (-) sense vRNA and (+) sense cRNA. Rather, cRNA and vRNA reads can be sorted based on their (+) and (-) polarity, respectively.

In addition to avoiding any of the previously discussed limitations of cDNA synthesis and PCR amplification strategies, the technique developed for direct RNA sequencing is highly amenable to sequencing a variety of non-poly-adenylated RNAs from hosts and pathogens, including untranslated regions (UTRs), without biasing the sequence to the primer. This allows the examination of the UTRs in their native form, which we have done here with influenza A virus. However, direct RNA sequencing of UTRs is limited by read level accuracy and a loss of coverage at the extreme 5’ end of the molecule. The extreme 3’ termini (Uni-12) of all segments were fully sequenced and matched the expected sequence with the exception of the degeneracy at the +4 position which was not resolved. The sequences for the extreme 5’ termini (Uni-13) that were obtained match the expected sequences with the exception of a C to G substitution at the -9 position in the segments PB1 and PB2. The loss of coverage at the extreme 5’ end of the molecule is most likely due to unreliable processivity as the last of the molecule passes and resulted in the final nine nucleotides not being sequenced in some of the segments.

The data presented demonstrates the adaptability of the platform and RNA sequencing protocol. The unmodified components were used to target enolase mRNA and could be used to target the variety of mRNA species present in any sample. Specifically, one could dissect viral replication processes as well as host mRNAs activated during an influenza infection at a given point in time. Genomic length and quantitative sequencing of viral mRNA species has the potential to provide direct detection of base modifications, splice variants, and transcriptional changes under different replication conditions, such as viruses used for vaccine production that are transferred between mammalian and avian hosts.

The primary limitations of this technology are the high read level error rate and high input material requirements. Reducing the error rate would enable multiplexing and more accurate consensus sequence determination and is a requirement for understanding nucleotide polymorphisms and genome sub-populations, particularly in viruses such as influenza that have significant intra-host diversity and or base modifications to be identified. There are currently several bioinformatic tools for detecting DNA base modifications such as Tombo, Nanopolish, SignalAlign, and mCaller; however, RNA specific tools have yet to be released^19^. Currently, the RNA input requirements for direct RNA sequencing are high and are not physically achievable with most original clinical samples. Lessening the RNA input requirement of the direct RNA sequencing would take full advantage of the unbiased nature of direct RNA sequencing and allow for the detection and description of the rich diversity intrinsic to influenza and other viruses. Although ONT has continuously improved their basecaller Albacore, there is still demonstrable potential for improvement. The RNA basecaller was likely developed using the very same enolase mRNA used here, which would make it most effective at basecalling enolase mRNA. The marked difference in accuracy between the enolase and influenza virus reads demonstrates that further development of the RNA basecaller can, at a minimum, bring the accuracy of all RNA reads up to that of enolase reads. Moreover, the DNA basecaller is overall more developed and more accurate than the RNA basecaller (89% versus 85% read level accuracy for influenza samples). The continued effort to advance this technology by ONT will undoubtedly result in higher accuracy reads and greatly improved utility.

## Methods

### Concentration and purification of A/Puerto Rico/8/1934 reassortant virus

A/Puerto Rico/8/1934 reassortant virus was grown in 11 day-old embryonated hen eggs at 35°C for 48 hours. Allantoic fluid was harvested from the chilled eggs and clarified at 5,400 × g, 10 minutes, 4°C, (Sorvall SLA-1500 rotor). The virus was clarified twice more by centrifugation at 15,000 × g, 5 minutes, 4°C (Sorvall SLA-1500 rotor). Virus was pelleted by centrifugation at 39,000 × g, 3 hours at 4°C (Sorvall A621 rotor). Virus pellets were resuspended overnight in PBS and loaded onto a 30%/55% (w/w) density sucrose gradient. The gradient was centrifuged at 90,000 × g for 14 hours at 4°C (Sorvall AH629 rotor). The virus fractions were harvested and sedimented at 131,000 × g (Sorvall AH629 rotor) for 2.5 hours. The resulting virus pellet was resuspended in PBS and aliquoted for future use.

### RNA isolation

Enolase II (YHR174W) mRNA is supplied in the ONT materials as the calibration RNA strand (CRS) at a concentration of 50 ng/μL. For influenza virus samples, total RNA was isolated by Invitrogen™ TRIzol^®^ extraction^21^ according to manufacturer’s instructions with additional considerations for biosafety. The virus was inactivated by the addition of 10 volumes of TRIzol^®^ in a Biosafely Level 2 biosafety cabinet. Following inactivation, a fume hood was used for the chloroform addition and aqueous phase removal steps. RNA pellets were resuspended in 10-40 μL nuclease free water and quantified by Quant-iT− RiboGreen^®^ RNA Assay Kit. Due to the difficulty in acquiring sucrose-purified material, the pure controls were limited to one MiSeq run and two separate MinION experiments. The availability of crude viral samples allowed it to be sequenced once on MiSeq and three times on MinION from the same RNA preparation.

### Nanopore Sequencing

The ONT direct RNA library preparation input material requirement is 500 ng of target molecule in a 9.5 μL volume (**Table S4**). For mRNA sequencing of the enolase control, the protocol was used according to the manufacturer’s instruction. For influenza viral RNA sequencing, modifications were made to the protocol components (**Table S2**). We altered the supplied reverse transcriptase adapter (RTA) which has a T10 overhang (Tm ~20°C) to target the ligation of the RTA to mRNA, with 12 nucleotides complementary to the conserved 3’ end of Influenza A virus (-) vRNA^22^ (**Figure 1**). RTA-U12 and RTA-U12.4 contained target sequences (5’ to 3’) AGC AAA AGC AGG and AGC GAA AGC AGG (Tm ~50°C) respectively and were combined in a 2:3 molar ratio to a total concentration of 1.4 μM. This mixture was used as a direct replacement to the RTA supplied in the protocol for influenza samples. Though there is some disagreement regarding the segment specific degeneracies of the 12 nucleotides at the 3’ end of the genome, RTA-U12 is expected to target the segments PA, NP, M, and NS; and RTA U-12.4 is expected to target the segments PB2, PB1, HA, and NA^23,24^.

Adapter ligated RNA was directly sequenced on the MinION nanopore sequencing using a FLO-MIN107 flowcell equipped with the R9.5 chemistry. The enolase sequencing experiments were operated through MinKNOW versions 1.4.2, 1.7.7, and 1.10.11; the pure sequencing experiments were operated through MinKNOW 1.7.7; and crude sequencing experiments were operated through MinKNOW 1.10.11. Raw data was basecalled using Albacore 2.1.10 (released 01/26/2018), and reads were assembled using IRMA^25^ with the FLU-MinION preset configuration to produce influenza consensus sequences for comparison to MiSeq-derived consensuses. The FLU-MinION preset differs from the default FLU module settings by the following: dropping the median read Q-score filter from 30 to 0, raising the minimum read length from 125 to 150, raising the frequency threshold for insertion and deletion refinement from 0.25 to 0.75 and 0.6 to 0.75 respectively, and lowering the Smith-Waterman mismatch penalty from 5 to 3 and the gap open penalty from 10 to 6. For read-level comparisons of MinION to MiSeq, raw fastqs from both sequencing platforms were mapped with bwa-mem v.0.7.7 algorithm^26^ to MiSeq+IRMA derived consensus sequences as references. Bwa-mem settings were left default except for the following arguments: “-A 2” and “-B 3”. Figures and tables were created in Tableau v.10.4.3.

Error rates were calculated against the aligned plurality consensus sequence as follows:

- Accuracy rate = 1 - average number of insertions, deletions, and minority alleles / sum of aligned bases + number of deletions and insertions at left-adjacent (upstream or 5’ to the site) base per position per segment
- Insertion rate = average number of insertions, irrespective of insertion length / sum of aligned bases + number of insertions at left-adjacent base per position per segment
- Deletion rate = average number of deletions, irrespective of deletion length / sum of aligned bases + number of deletions at left-adjacent base per position per segment
- Substitution rate = average number of minority bases / sum of aligned bases per position per segment.
- Alignment read lengths were calculated as matching + inserted bases per read (CIGAR M+I).

### Illumina MiSeq Sequencing

The complete influenza genome was amplified with the RNA from both the sucrose purified virus and the allantoic fluid. The MRT-PCR used the Uni/Inf primer set^16^ with SuperScript III One-Step RT-PCR with Platinum Taq High Fidelity (Invitrogen). Following amplification, indexed paired-end libraries were generated from 2.5 μl of 0.2 ng/μL using the Nextera XT Sample Preparation Kit (Illumina) following the manufacturer protocol using half-volume tagmentation reactions. Libraries were purified with 0.8X AMPure XP beads (Beckman Coulter, Inc.) and assessed for fragment size (QIAxcel Advanced System, Qiagen) and quantitated using Quant-iT dsDNA High Sensitivity Assay (Invitrogen). Six pmol of pooled libraries were sequenced on the Illumina MiSeq with MiSeq v2 300 cycle kit and 5% PhiX spike-in to increase the sequence diversity. Sequence analysis was performed using IRMA^25^ as part of the current Illumina-based pipeline utilized by the Influenza Genomics Team at the Centers for Disease Control and Prevention.

## Acknowledgements

Research reported in this publication was supported by the office of Advanced Molecular Detection (AMD CAN 939018C) at the Centers for Disease Control and Prevention. We thank Oxford Nanopore Technology’s technical support team, Bryant Catano in particular, for the recovery of QC data from early sequencing experiments.

## Author contributions statement

D.W. and J.B. conceived the research. M.K., M.W., and C.R. conducted the experiments. M.K., B.R-M., T.S. and S.S. analyzed the results. B.R-M. accessioned the raw data. M.K., B.R-M., M.W., C.R., S.S., T.S., E.N., V.D., D.W. and J.B. edited the manuscript.

## Competing financial interest

The authors declare no competing financial interests.

## Data availability

Sequence data is accessioned at NCBI: <u>SRP139094</u>.

## Supplementary Information

**Figure S1.**
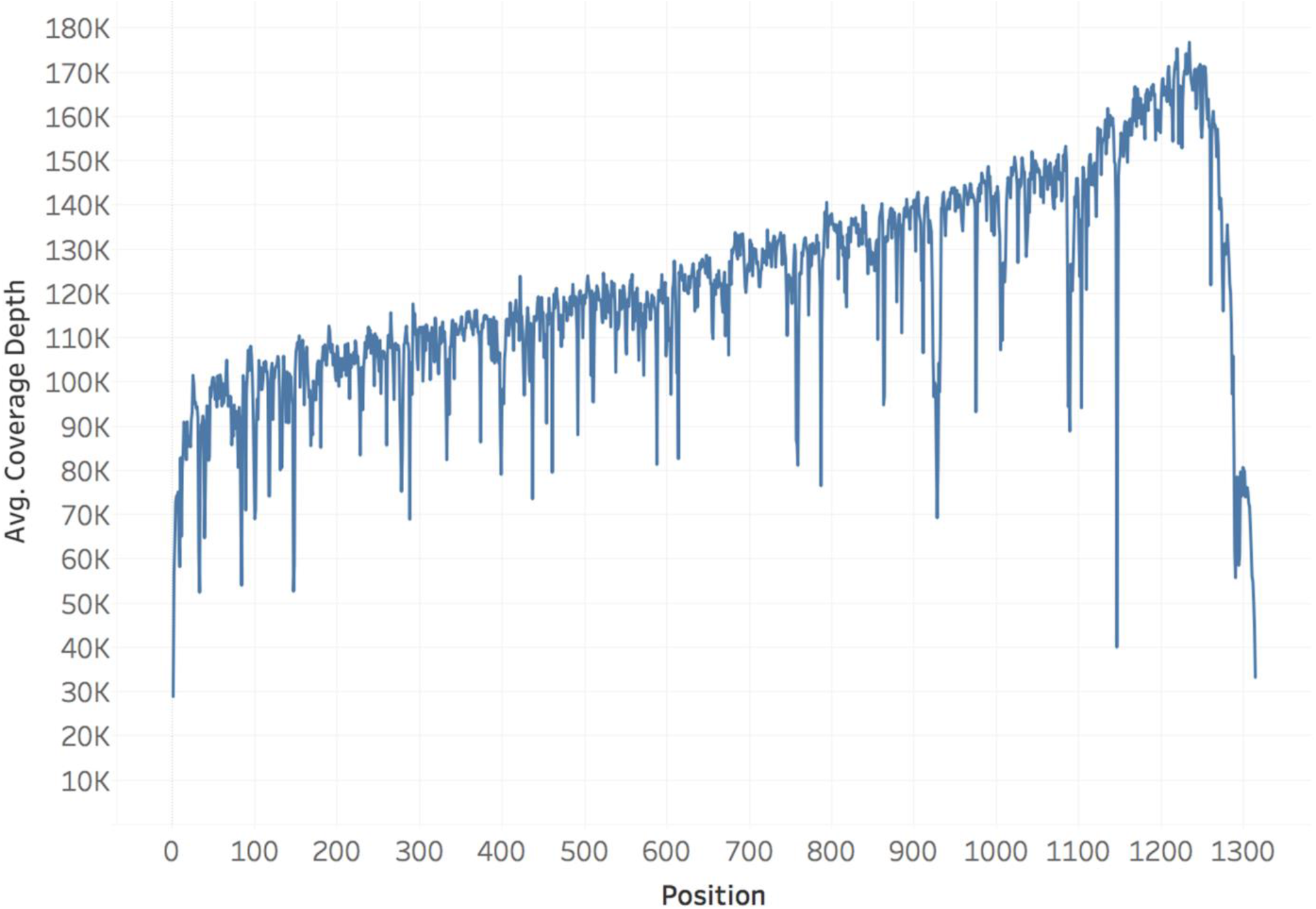
Average coverage of triplicate enolase direct RNA sequencing experiments. The MinION was able to sequence enolase mRNA to an average coverage depth of 117,408 ± 9,617 reads. The directional nature of nanopore sequencing results in a positive slope to the coverage for the mRNA.

**Figure S2.**
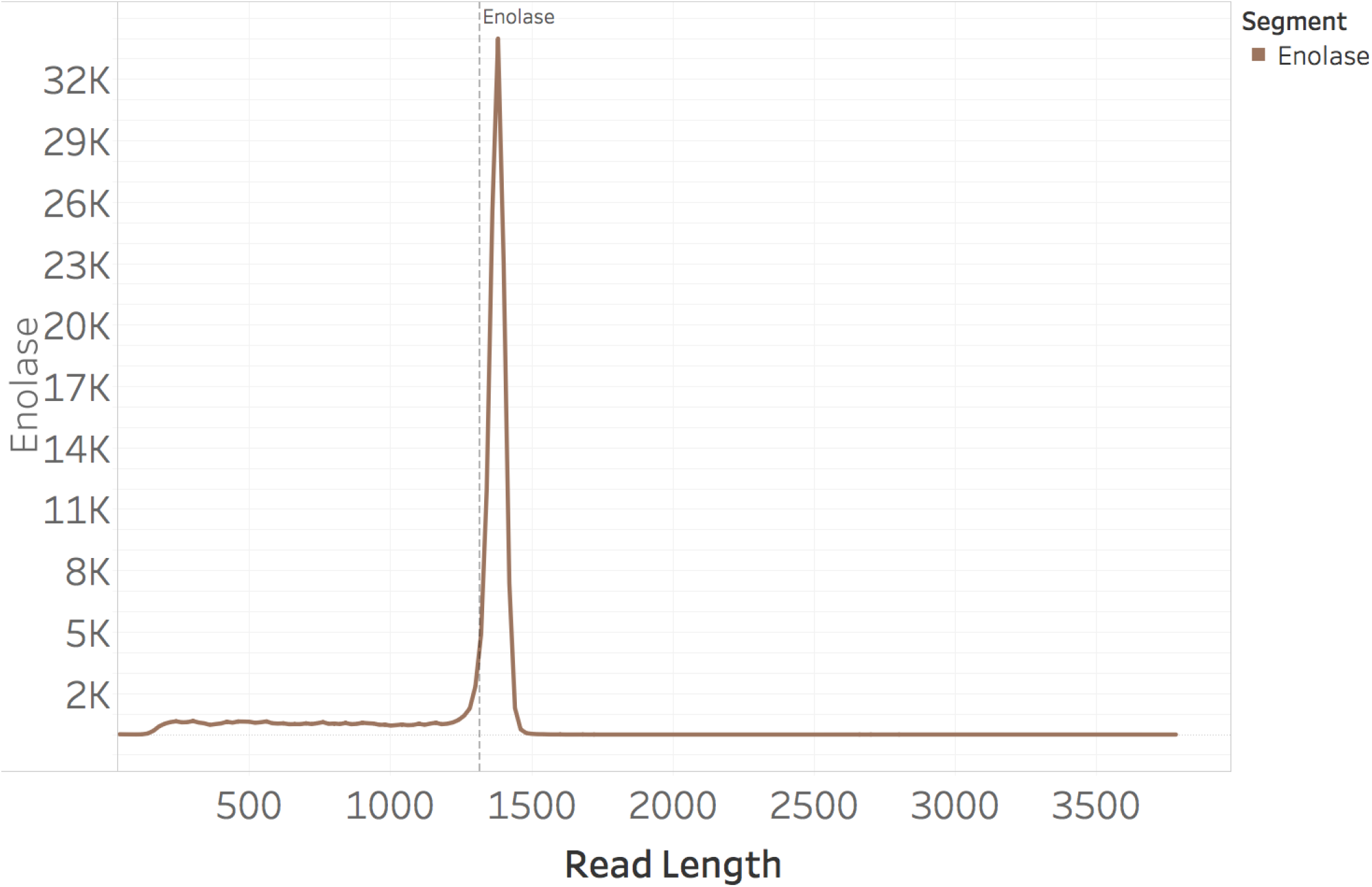
The aligned read length distribution is longer than the expected length of 1,314 nucleotides (dashed line) due to insertions.

**Figure S3.**
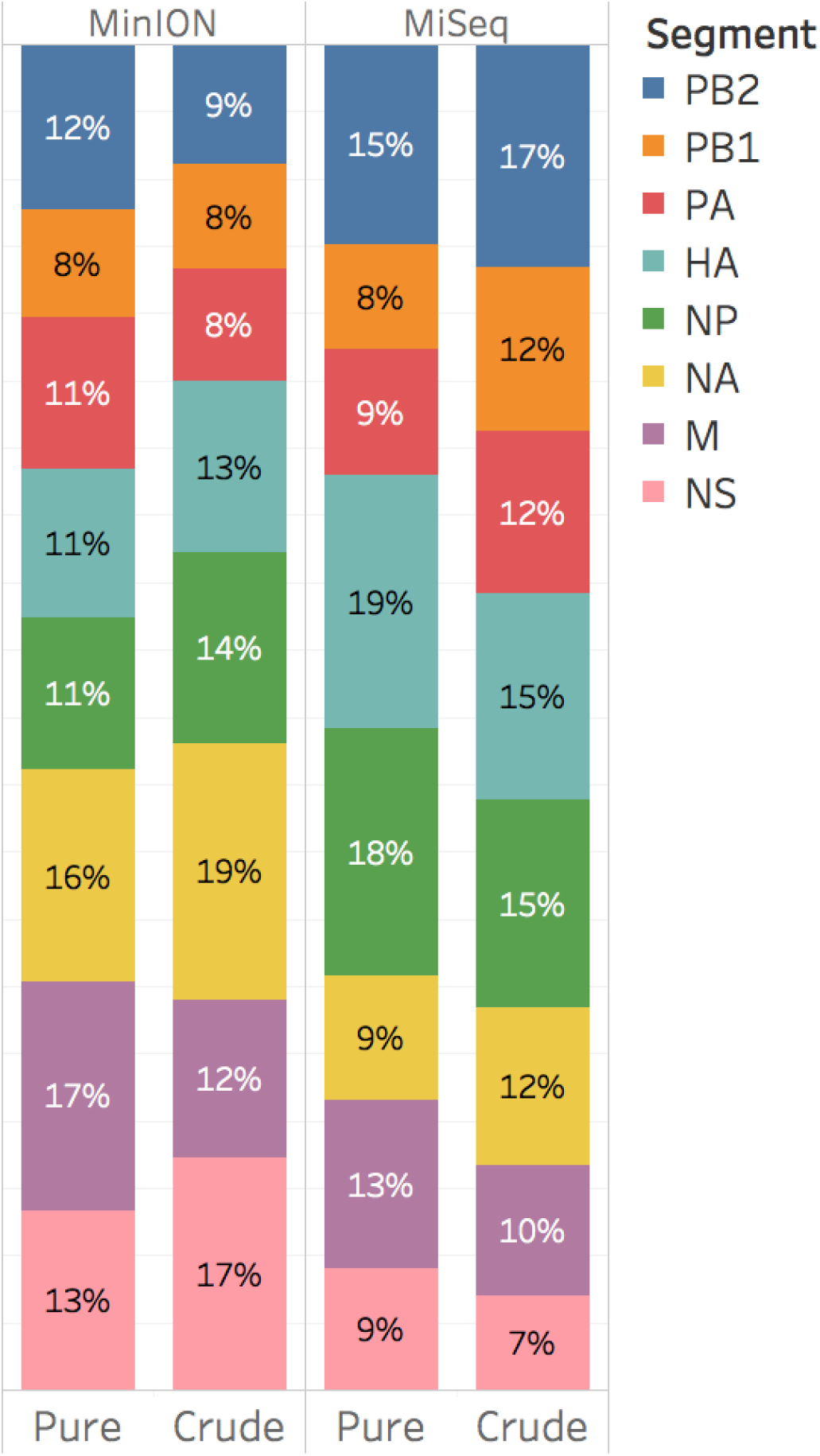
Distribution of mapped reads for all experiment iterations.

**Figure S4.**
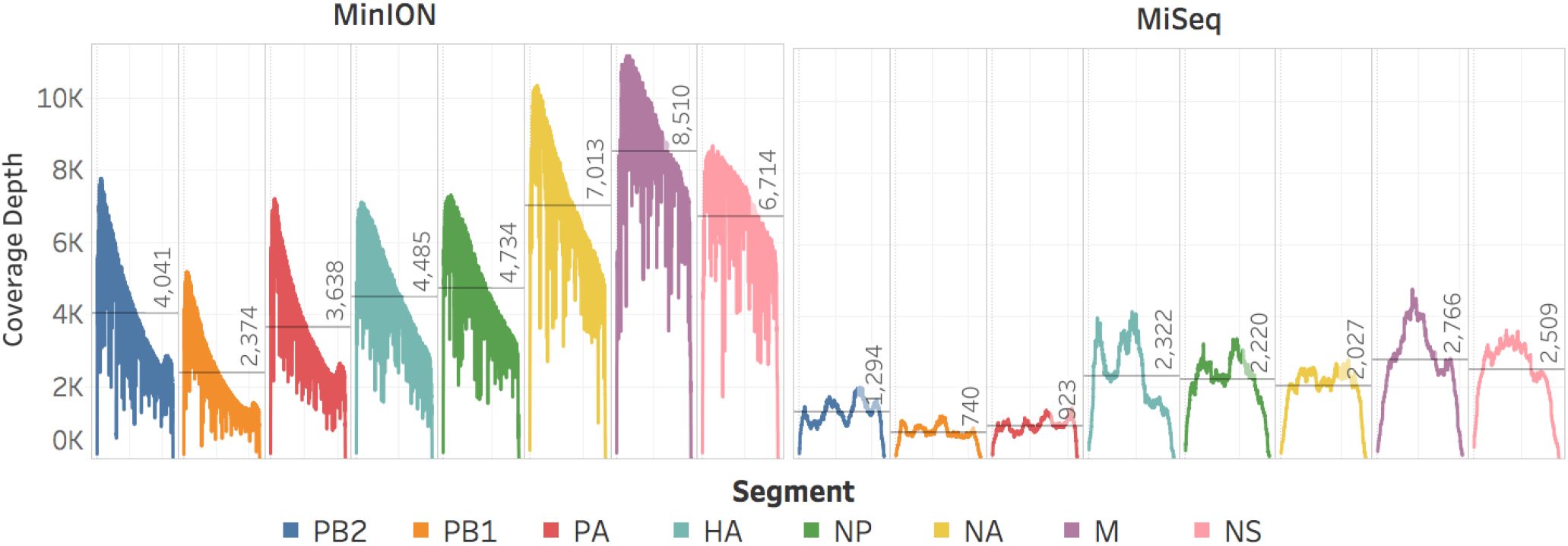
MinION and MiSeq sequencing coverage of the PB2, PB1, PA, HA, NP, NA, M, and NS genome segments of the pure influenza virus genome. Negative-sense slope coverages in the MinION results confirm the directionality of the sequencing and capture method.

**Figure S5.**
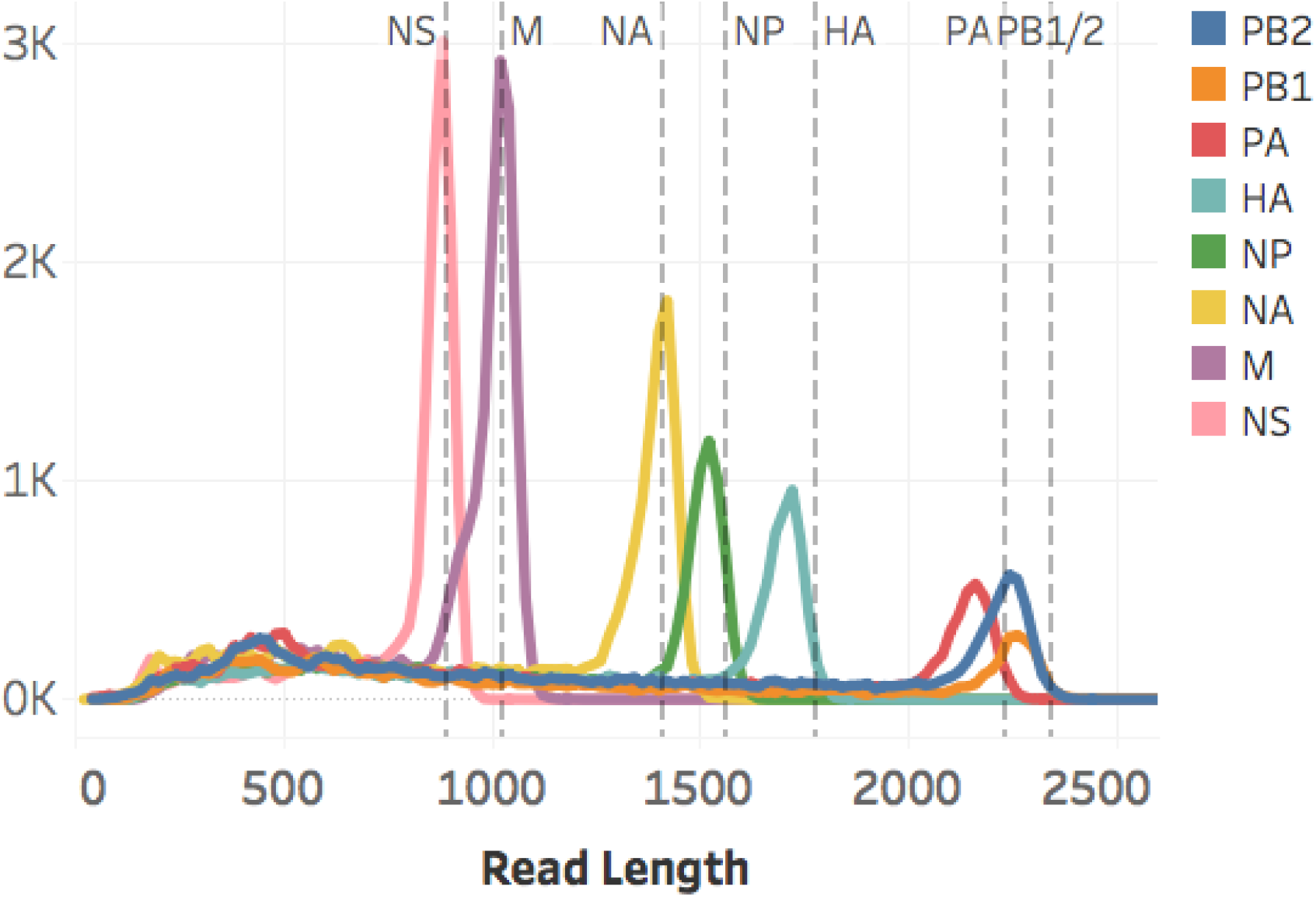
The aligned read length distributions correspond to the expected lengths (dashed lines) of the respective segments (NS 890 nt; M 1,027 nt; NA 1,413 nt; NP 1,565 nt; HA 1,778 nt; PA 2,233 nt; PB1 and PB2 2,341 nt). As the segment length increases, the read length distribution falls further short of the expected length, presumably due to RNA degradation. Aligned read lengths include insertion errors, accounting for the presence of reads larger than the expected value. Due to cases of large insertion errors, 14 total reads longer than 2,500 nucleotides were observed.

**Figure S6.**
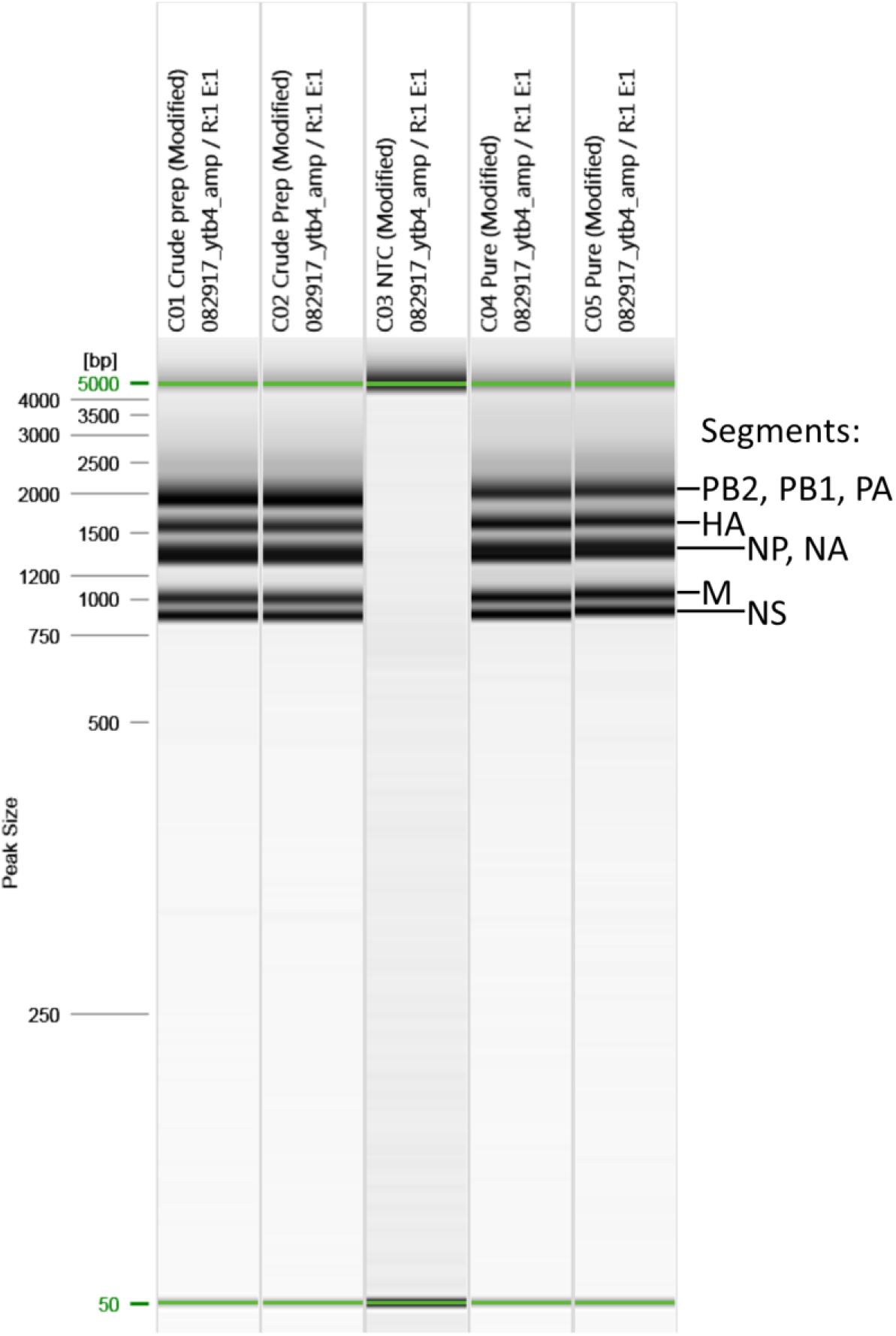
QIAxcel size fractionation and visualization of amplicons from (left to right): crude duplicates, negative control, and pure duplicates. The five visible bands in lanes 1, 2, 4, and 5 are characteristic of uniform amplification of the entire influenza virus genome. The PB2, PB1, PA, HA, NP, NA, M, and NS genome segments are represented by only five bands because the three segments that encode the polymerases appear as a single band just larger than 2,000 and the NP and NA segments appear as a single band just below 1500.

**Figure S7.**
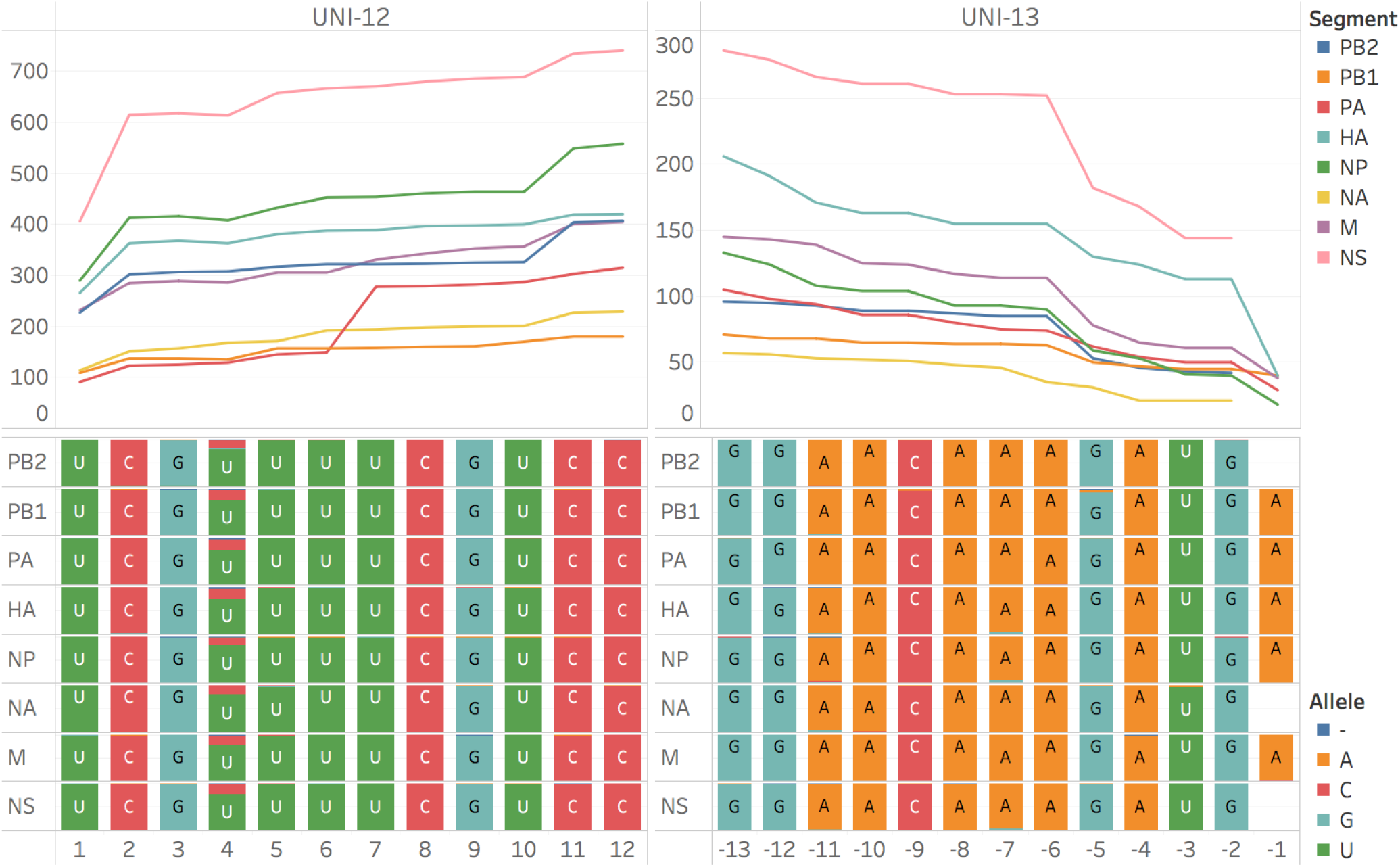
MiSeq coverage and consensus of the conserved termini of the influenza viral RNA genome segments from the crude viral samples. These are the amplification sites, and the results are primer dictated sequences.

**Table S1.** Average and mode mapped read length is shown for MinION direct RNA sequencing experiments. The presence of short reads, particularly in the crude sample, move the average read length much lower than the mode read length that is displayed here and in **figures 4, S2, and S5**. The read length distribution of the polymerases are all bimodal with an abundance of short reads along with full length reads. The read length distribution of M from the crude viral sample is also bimodal with a clear and well-defined peak at 920 nucleotides in addition to the full-length peak at 1,020 nucleotides. All Illumina reads were 150 nucleotides in length.

| Gene & Length |  | Average Length |  | Mode Length |  |
| --- | --- | --- | --- | --- | --- |
|  |  | Enolase/Pure | Crude | Enolase/Pure | Crude |
| Enolase | 1,314 | 975 | - | 1380 | - |
| PB2 | 2,341 | 1,177 | 752 | 440/2,240 | 280/2,240 |
| PB1 | 2,341 | 1,061 | 718 | 580/2,260 | 260/2,260 |
| PA | 2,233 | 1,089 | 833 | 500/2,160 | 280/2,140 |
| HA | 1,778 | 1,111 | 828 | 1,720 | 1,720 |
| NP | 1,565 | 1,025 | 741 | 1,520 | 1,520 |
| NA | 1,413 | 980 | 711 | 1,420 | 1,400 |
| M | 1,027 | 809 | 649 | 1,020 | 920/1,020 |
| NS | 890 | 698 | 607 | 880 | 880 |

**Table S2.**
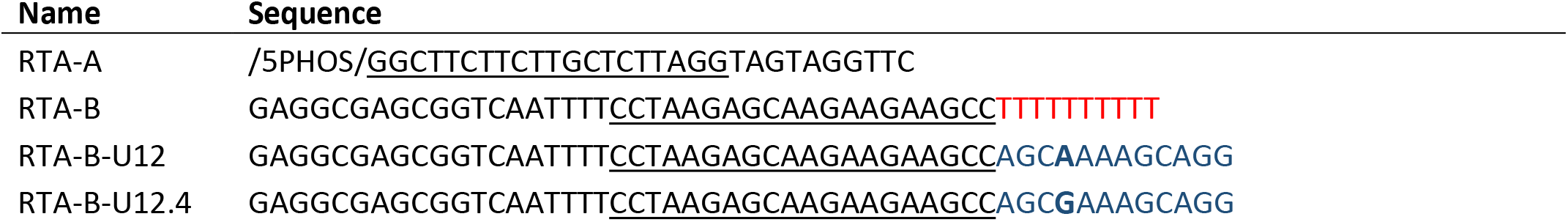
Full sequences (5’ to 3’) of the adapters used in this study. Each RTA-B is duplexed with RTA-A. The stock RTA was supplied with the direct RNA sequencing materials. The modified RTAs were purchased from IDT with each of the modified RTA-B strands already duplexed to the RTA-A strand. The RTA-A has a 5’ phosphate modification for ligation. The regions of reverse complementarity between the RTA strands are underlined, and the target sequences are colored.

**Table S3.**
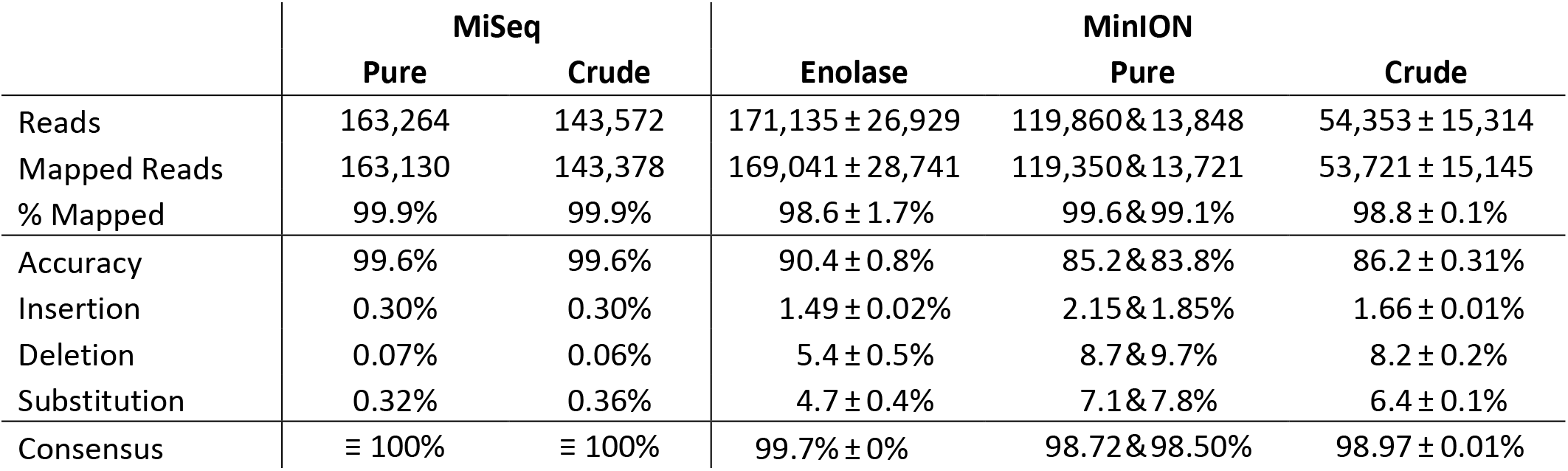
Individual MiSeq experiments are compared to MinION experiments of enolase mRNA (technical triplicate), influenza vRNA from purified virus (duplicate), and influenza vRNA from crude virus (triplicate). Values from triplicate experiments are presented as averages ± standard deviation.

**Table S4.** Input material and sequencing information for the direct RNA sequencing experiments. Pore availability for mux 1-3 is displayed. These data could be recovered by ONT technical support for Enolase 1 or Pure 1. Mux 4 scan data was not able to be recovered for these samples. The full mux (1/2/3/4) data for the other samples was: enolase 2 (420/202/44/12), enolase 3 (509/492/403/180), pure 2 (291/110/13/1), crude 1 (483/390/229/58), crude 2 (506/456/327/134), and crude 3 (449/393/230/53). The mux 4 pores are fewer in number and are used lastly in long sequencing experiments, if they are used at all. Pore occupancy was estimated by ONT technical support from QC reports.

| Date | Sample | RNA (ng) | Library (ng) | Flowcell | Pores | Occupancy | Time (hours) | Yield (Mb) |
| --- | --- | --- | --- | --- | --- | --- | --- | --- |
| 2017-06-19 | Enolase 1 | 500 | 180 | FAH04399 | 745 | 35% | 48 | 180 |
| 2017-08-04 | Enolase 2 | 500 | 133 | FAH14471 | 666 | 21% | 48 | 197 |
| 2017-11-09 | Enolase 3 | 500 | 32 | FAH28081 | 1,404 | 27% | 48 | 172 |
| 2017-07-13 | Pure 1 | 216 | 60 | FAB49814 | 656 | 45% | 24 | 127 |
| 2017-09-08 | Pure 2 | 257 | 254 | FAH15505 | 414 | 5% | 14 | 14 |
| 2017-11-22 | Crude 1 | 200 | 18 | FAH36033 | 1,102 | 18% | 18 | 59 |
| 2017-11-27 | Crude 2 | 200 | 15 | FAH28243 | 1,289 | 10% | 16 | 35 |
| 2017-11-28 | Crude 3 | 200 | 15 | FAH36048 | 1,072 | 10% | 17 | 34 |

